# The Hox Gene, *abdominal-A*, controls the size and timely mitotic entry of neural stem cells during CNS patterning in *Drosophila*

**DOI:** 10.1101/2024.09.04.611161

**Authors:** Papri Das, Smrithi Murthy, Eshan Abbas, Kristin White, Richa Arya

## Abstract

Cell size is a critical determinant of its function and physiology. In this study, we investigated the regulation of stem cell size during *Drosophila* central nervous system (CNS) development and its association with cell fate. We note that neural stem cells (NSCs) in different regions of ventral nerve cord increase their size at different rates. The, thoracic NSCs grow at a faster rate compared to those in the abdominal region. We show that in addition to its role in apoptosis and nervous system remodeling, *abdA* also plays an instructive role in regulating the rate of NSC size increase and its timely entry into mitosis. Weak expression of abdA in larval abdominal NSCs was sufficient to retard the rate of their size increase and delay their mitotic entry compared to thoracic NSCs. Knockdown of *abdA* in NSCs enhances their rate of size increase leading to early mitotic entry, while ectopic *abdA* expression in thoracic NSCs reduces their size and delays their mitotic entry. We show that *abdA*-mediated NSC size regulation acts downstream of nutrition-induced NSC activation, which fine-tunes the stem cell potential spatiotemporally. This study highlights the instructive role of *abdA* in regulating various fates of larval NSCs during CNS patterning.

**Significance statement:**

- Understanding the upstream regulation of various aspects of the cell cycle is crucial; however, the influence of cell size on this process remains largely unknown.
- We found an instructive role for the Hox gene *abdominal A* in maintaining the small size of neural stem cells (NSCs) and in regulating the time and rate of mitosis.
- This mechanism is crucial because it helps NSCs generate the necessary number of neurons at an appropriate developmental stage, thereby contributing to proper central nervous system patterning.

## Introduction

Multipotent neural stem cells (NSCs), possess the capacity to generate a diverse range of cell types. The conservation of molecular signaling involved in NSC fate determination, combined with the availability of genetic tools for studying gene functions, makes *Drosophila* an excellent model for exploring stem cell fate determination at the molecular level (Allan and Thor, 2015; Doe, 1992; Homem and Knoblich, 2012; Livesey and Cepko, 2001; Verma et al., 2019). During early embryonic development, NSCs delaminate in a segment-specific manner, dividing asymmetrically to self-renew and produce neurons and glial cells (Hartenstein et al., 1994; Truman and Bate, 1988). Interestingly, the timing of larval neurogenesis onset varies by region, with NSCs in the anterior region typically beginning to proliferate earlier than those in the posterior region (Britton and Edgar, 1998; Prokop and Technau, 1994; Prokop et al., 1998; Truman and Bate, 1988).

Changes in cell size are key outcomes of cellular growth and play an essential role in regulating various aspects of a cell’s function and fate. The metabolic status, specialized secretion, transport, proliferation, and many other cellular activities depend on cell size (Cadart et al., 2018; Ginzberg et al., 2018; Kafri et al., 2013; Miettinen and Björklund, 2016). In *Drosophila*, neural stem cells (NSCs) exhibit dynamic size changes during neurogenesis, increasing in size at the onset and decreasing at the end of this process. After embryonic neurogenesis, NSCs in the ventral nerve cord (VNC) become smaller before entering a quiescent state. At the onset of larval life, these cells gradually enlarge before entering mitosis (Britton and Edgar, 1998; Colombani et al., 2003; Hartenstein et al., 1987a; Prokop and Technau, 1991; Truman and Bate, 1988). There is size heterogeneity among NSCs in the larval VNC, with differences observed between the thoracic and abdominal segments. Notably, mitotic NSCs in the thoracic region are consistently larger and divide faster than those in the abdominal region (Truman and Bate, 1988). Similarly, *Drosophila* midgut stem cells display differences along their anterior-posterior axis, maintaining regional identity reflected in their morphology, gene expression, and proliferation rates (Marianes and Spradling, 2013). In mammalian systems, the growth and maintenance of hematopoietic stem cell (HSC) size are also critical for cell function (Lengefeld et al., 2021; Saçma et al., 2025). In young and aged mice, HSCs exhibit distinct size differences across various niches. Smaller HSCs tend to be more myeloid-biased, while larger HSCs show a preference for B-lymphoid differentiation (Saçma et al., 2025). In contrast to *Drosophila* NSCs, proliferating HSCs are generally smaller, and their proliferative potential significantly decreases as their size increases due to aging (Lengefeld et al., 2021). Overall, it appears that stem cell size plays a crucial role in determining cell behavior. Although well-known pathways regulating cell growth, such as the Hippo, Insulin, and PI3K pathways, influence NSC size, (Chell and Brand, 2010; Ding et al., 2016; Elkin et al., 2025; Gil-Ranedo et al., 2019; Sousa-Nunes et al., 2011), the mechanisms that maintain size heterogeneity remain unclear. Investigating the spatiotemporal regulators of NSC growth rates and their connections to cell fate and function would provide insight into both development and disease.

Homeotic (Hox) genes play a crucial role in development and tissue patterning, primarily by establishing segment-specific identity and function. (Bender et al., 1985; González-Reyes et al., 1992; Jiménez and Campos-Ortega, 1981; Lewis, 1978; Mann and Morata, 2000; Sánchez-Herrero et al., 1985; Wakimoto and Kaufman, 1981). *Abdominal-A* (*abdA*), a member of the bithorax complex in *Drosophila*, is expressed in the abdominal region of the developing nervous system along the anterior-posterior axis and play a crucial role in determining the abdominal identity of neural lineages (Bender et al., 1985; Lewis, 1978; Prokop and Technau, 1994b; Sánchez-Herrero et al., 1985). In addition to providing spatial identity to abdominal NSCs, *abdA* regulates their temporal fate by limiting the time of abdominal NSC proliferation and terminal fate by controlling their removal from the nervous system by apoptosis (Arya et al., 2015; Bello et al., 2003; Prokop et al., 1998). Under the influence of *abdA*, all abdominal NSCs undergo apoptosis in two distinct waves: the first occurs at the end of embryonic development, and the second takes place during mid-larval life (Abrams et al., 1993; Peterson et al., 2002; White et al., 1994, Bello et al,. 2003).Interestingly, despite evidence that *abdA* regulates the proliferation of larval abdominal NSCs, its expression has been reported only during early embryonic stages, leading to the suggestion that it is unlikely to be a direct regulator of their proliferation (Prokop et al., 1998). Furthermore, the specific factors that limit the growth of abdominal NSCs remain unknown.

In this study, we found that from the onset of larval neurogenesis, both thoracic and abdominal NSCs in the *Drosophila* VNC gradually increased in size, although at different rates. Our results indicated that the rate of size increase for the abdominal NSCs was slower compared to those in the thoracic region. The size of NSCs is a critical factor influencing their ability to initiate proliferation (Britton and Edgar, 1998; Chell and Brand, 2010; Truman and Bate, 1988; Yuan et al., 2020). Although abdominal NSCs enter the S phase at the same time as thoracic NSCs, they experience a prolonged G2 phase and enter mitosis much later and divide at a slow rate during larval development, making fewer progeny than thoracic NSCs (Prokop et al., 1998; Truman and Bate, 1988). We show that when *abdA* expression is lost, abdominal NSCs increase in size at a faster rate, have a shorter G2 phase, enter mitosis earlier, and divide at a faster rate. Conversely, the gain of *abdA* in thoracic NSCs can slow their growth and delay their entry into mitosis. In addition to the known role of *abdA* in regulating developmental apoptosis of abdominal NSCs (Arya and White, 2015; Bello et al., 2003), this study highlights *abdA*’s role in controlling the size, mitosis, and proliferation rates of NSCs during *Drosophila* CNS patterning.

## Results

### The NSCs in larval *Drosophila* VNC show a spatially restricted size distribution between the thoracic and abdominal regions

To understand the time at which size differences emerged, we profiled the growth patterns of NSCs in the VNC at embryonic to various larval time points (Fig. 1A-G’, I, J). We used the HES1 protein analog Deadpan (Dpn) to mark the NSC nuclei (Fig.1K) on VNC of CNS, and membrane GFP driven with an NSC-specific *Gal4* driver inscutable (*insc*) (*insc>mCD8-GFP*) to mark the NSC cell membrane. Abdominal and thoracic NSCs are reported to be of different sizes in larval VNC (Truman and Bate, 1988). All embryonic NSCs in the VNC undergo quiescence from stage 15 onwards and exit quiescence during larval life when nutrition is available (Britton and Edgar, 1998; Chell and Brand, 2010; Sousa-Nunes et al., 2011). We measured the size of embryonic thoracic and abdominal NSCs during stages 16 and 17, when most NSCs were quiescent. We found no significant difference in the size of thoracic and abdominal NSCs (mean size 4.11, 4.2 µm of thoracic and abdominal NSCs respectively), indicating that all NSCs in the embryonic VNC are at a similar size during the late stages of embryonic quiescence. (Fig. 1I,J; Supplementary Fig. 1A,B Fig. 2B).

**Figure 1.**
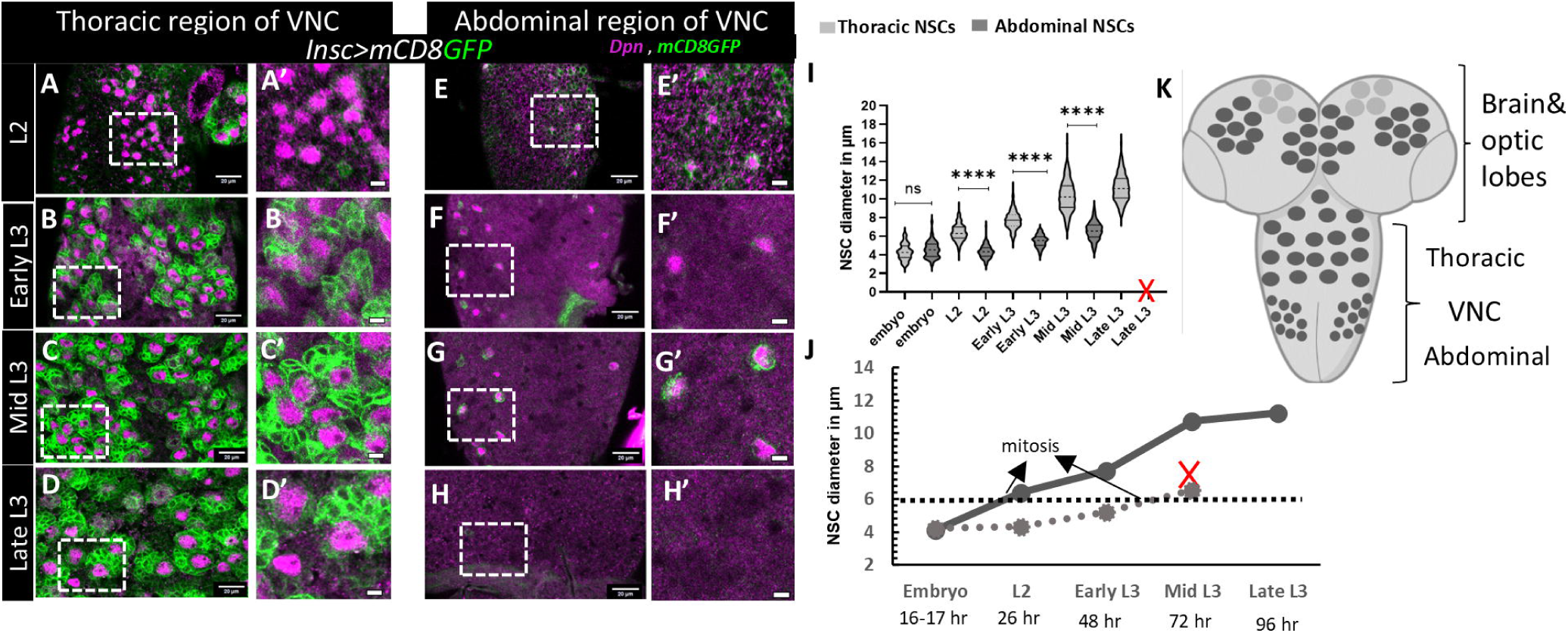
The NSCs in larval *Drosophila* VNC show spatially restricted size distribution between thoracic and abdominal regions. (A-D) thoracic NSCs (A’-D’) are magnified areas of the same images. (E-H) abdominal NSCs (E’-H’) are magnified areas of the same images. In all images, NSC nuclei are marked by Deadpan (magenta) and the membrane is marked by mCD8GFP (green), which also marks their progeny (*insc>UAS-mCD8GFP*). (I) Graphically depicts NSC size in the thoracic and abdominal regions during different developmental stages. “X” in the graph indicates developmental apoptosis of NSCs. (J) shows how thoracic and abdominal NSCs gradually increase in size during different developmental stages (K) A model depicting *Drosophila* CNS with optic lobes and VNC, divided into thoracic and abdominal regions, showing type 1 NSCs (grey circles) and bigger Mushroom body NSCs (light grey circles). Statistical evaluation of significance based on an unpaired t-test is marked with asterisks; ****P<0.0001. Scale bar: 20 µm and 5 µm for magnified images. More than five CNS were analyzed for each case.

**Figure 2.**
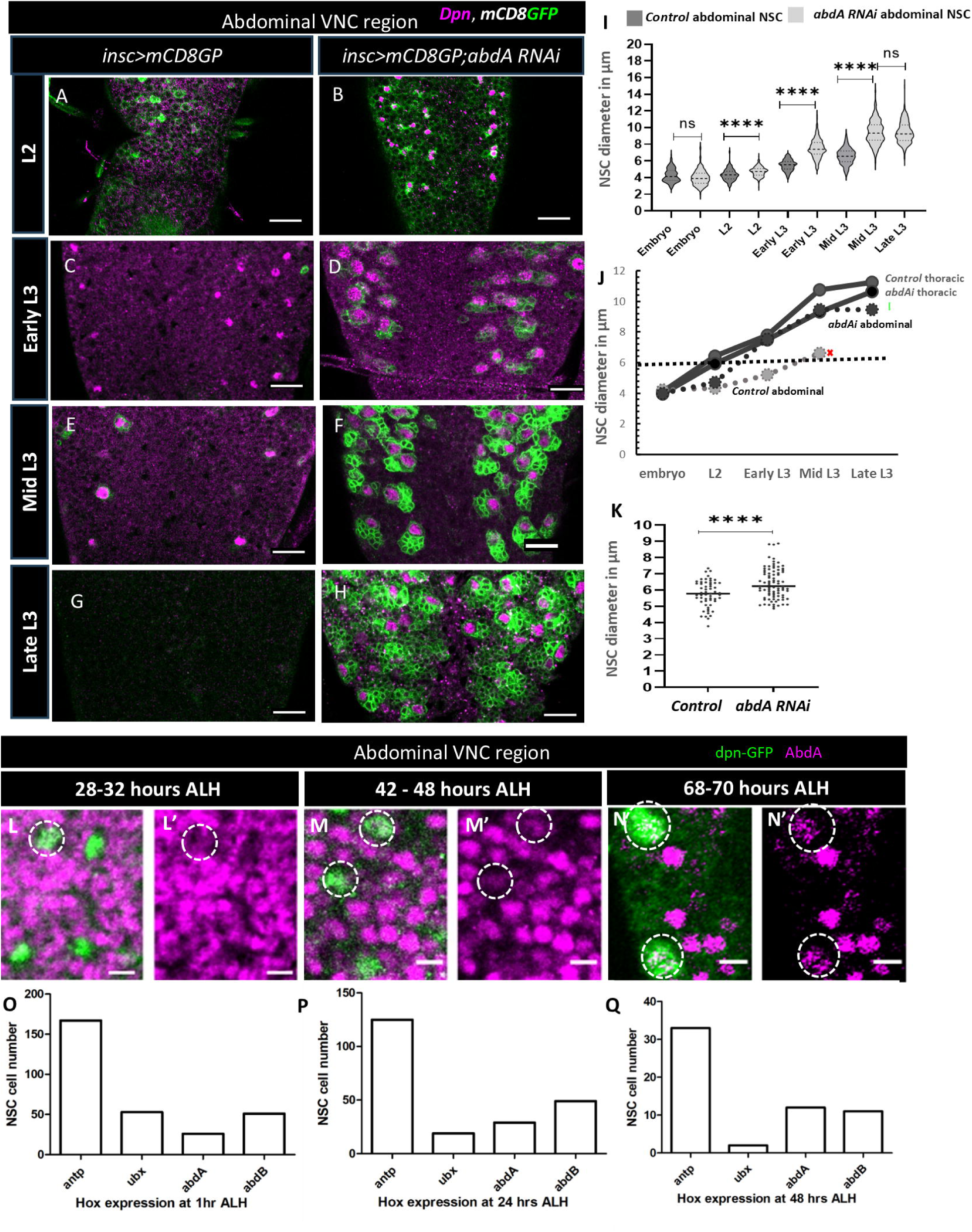
*abdA* restricts the rate of abdominal NSCs size increase during larval development. (A–H) Abdominal NSCs increased their size over the larval stages (from L2 to late L3), (A, C, E,G *insc>mCD8GP*) controls where (G) shows no NSCs, due to developmental apoptosis; (B,D,F,H *insc>mCD8GP;abdA-RNAi*) *abdA* knockdown showing pronounced increase in NSCs size (and number due to rescue from apoptosis) at different larval stages. (I-J) Graph showing the increase of abdominal NSCs size in control and *abdA* knockdown from L2 to Late L3 stages. “X” indicates developmental loss of NSCs in late L3 due to apoptosis. (K) abdominal NSC size at Mid L3 in both control and *abdA* knockdown when *worniu-Gal4* was restricted to express from early larval life with *Gal80ts* (control *Gal80^ts^::Wor-Gal4/+ and abdA knockdown Gal80^ts^::Wor-Gal4> abdA-RNAi*). (L-N’) Single sections of confocal images showing abdA expression (magenta) in abdominal region during different larval stages, NSCs (marked with dpn-GFP, and abdA are outlined with white dotted circles). (O-Q) Bar plots of scRNA-Seq showing the number of NSCs that express the four Hox genes, *Antp*, *Ubx*, abdA and *AbdB* in the larval VNC at 1 hr, 24 hr, 48 hr ALH(After Larval Hatching), respectively (Dillon et al. 2022). An expression threshold (E) > = 0.5 is considered positive. The expression threshold was set for logarithm-normalized expression levels. Statistical evaluation of significance based on an unpaired t-test is marked with asterisks; ****P<0.0001. Scale bar: 20 µm. More than five CNS were analyzed in each case.

We found that during the larval stages, the nuclei of abdominal NSCs were smaller than those in the thoracic region (Fig. 1A-G’). Since the size of the nucleus is generally proportional to the cytoplasmic volume across different cell types, we also checked the cell size of NSCs. We observed the same spatially restricted cell size patterning when the membranes of each NSC were stained with GFP (Fig. 1A-G’, I, J). During the larval stages, these NSCs of thoracic and abdominal region differentially increased their size and maintained a size ratio of thoracic vs abdominal NSC with a mean value of approximately 1.5, during larval stage 2 (L2), early larval stage 3 (early L3), and mid-larval stage 3 (Mid L3) (Fig. 1I,J; Supplementary Fig. 2B). This indicated that the thoracic NSCs were larger than the abdominal NSCs during all larval stages. The size difference between abdominal and thoracic NSCs at late larval stage 3 (LL3) could not be ascertained because at this stage, all abdominal NSCs were eliminated by apoptosis as previously mentioned (Bello et al., 2003) (Fig. 1H-H’). Here, we show that the pattern of NSC cell size increase in VNC remains spatially restricted in thoracic and abdominal regions throughout larval life.

### Hox gene *abdA* restricts the rate of abdominal NSCs size increase during larval development

To investigate the role of spatial transcription factors in regulating the size of NSCs, we examined the Hox gene *abdA*, which exhibits restricted expression in the central abdominal region of the VNC (Lewis, 1978; Sánchez-Herrero et al., 1985, Supplementary Fig. 2A). *abdA* is crucial for defining the spatial identity of the abdominal VNC region and is expressed in several neural cells starting from the embryonic stage. (Arya et al., 2015; Bender et al., 1985; Lewis, 1978; Prokop et al., 1998; Sánchez-Herrero et al., 1985). Basal expression of abdA in NSCs can be seen during stage 10-11 embryos, and it is also required for their timely elimination by apoptosis during late embryonic life (Arya et al., 2015; Prokop et al., 1998, Lewis, 1978). Temporal induction of abdA during late larval life correlates with the removal of the remaining abdominal NSCs through apoptosis (Bello et al., 2003).

To understand the role of *abdA* in regulating NSC size, we knocked it down using RNAi in NSCs starting from embryonic life. We evaluated NSC size at different larval stages, using the membrane marker mcd8-GFP (*insc>mCD8GFP,abdA-RNAi*). After the *abdA* knockdown, the number of abdominal NSCs significantly increased at larval stage L2, due to inhibition of apoptosis during embryonic development as previously reported (Aryaet al.,et al.,2015, Supplementary Fig. 2M). We observed a statistically significant difference in the sizes of the control (mean size 4.3μm) and *abdA*-depleted abdominal NSCs (mean size 4.7μm) at this stage (Fig. 2A,B,I,J). By early L3, a notable size difference between abdominal NSCs in *abdA*-knockdown VNCs and controls was evident (mean size in control 5.19μm, *abdA-*depleted NSCs 7.5μm), showing that the *abdA*-depleted NSCs were considerably larger than those in the control group (Fig. 2C-H,I,J). This steady increase in the size of *abdA*-depleted NSCs from stages L2 to L3 was consistent compared to control NSCs (Fig. 2I,J). Since abdA expression is limited to the central abdomen, the size of the thoracic NSCs remained unaffected (Fig. 2J and Supplementary Fig. 2 D,E). In the late embryonic stages 17 (image not shown), we also compared the size of abdominal NSCs in the VNC of control and *abdA* knockdown conditions and found no significant difference (mean size 4.2μm for control and 4.0μm for *abdA*-depleted NSCs) (Fig. 2J). These findings suggest that the depletion of *abdA* does not influence the size reduction of embryonic NSCs and its role is more restricted to the larval development.

To further investigate whether the effect of abdA loss on NSC size is confined to its expression during larval life, we restricted Gal4 expression to larvae using temperature-sensitive Gal80 (*Gal80^ts^::Wor-Gal4>abdA-RNAi,* Supplementary Fig. 2C,F,G). Indeed, we observed a significant increase in the size of *abdA-*depleted larval NSCs compared to controls, indicating that *abdA* restricts larval NSC size (Supplementary Fig. 2C).

We compared the sizes of abdominal and thoracic NSCs in both the control and the *abdA* knockdown groups. Our observations indicated a reduced size difference between thoracic and abdominal NSCs following *abdA* knockdown. Typically, thoracic NSCs are 0.4 to 0.6 times larger than abdominal NSCs at various larval development stages. However, this size difference decreased to a range of 0.0 to 0.3 when *abdA* was depleted from the abdominal NSCs (Fig. 2J; Supplementary Fig. 2B,D,E). Considering that Hox genes often adhere to the posterior prevalence rule, we also investigated whether *abdA* knockdown caused ectopic expression of Ubx in the abdominal NSCs, which normally express in the region just prior to the abdA expression domain (Bender et al., 1985; Lewis, 1978; Sánchez-Herrero et al., 1985),. We found no expression of Ubx in the rescued NSCs (Supplementary Fig. 2L-Y), indicating that abdA expression in abdominal NSCs plays a role in regulating their size.

Previous research has indicated that abdA is only expressed in abdominal NSCs as a pulse between 64-72 hours after larval hatching (ALH) to initiate apoptosis (Bello et al., 2003). Since our results indicated a significant effect of abdA on the size of abdominal NSCs beginning in early larvae, we undertook a detailed description of the expression pattern of abdA in the NSCs during the larval period from L1 to L3 (68-70 hr ALH), before the time of abdominal NSCs death (fig. 2L-N’). We used a MiMIC line of *dpn-GFP*, in which endogenous *dpn* was tagged with GFP (Nagarkar-Jaiswal et al., 2015). During the very early larval stage (14-17 hr ALH), dpn-GFP was not detectable in any of the thoracic NSCs. Abdominal NSCs also showed low or no dpn expression during this stage. abdA showed widespread differential expression in many cells in the abdominal region, with some cells expressing more abdA than others (Supplementary Fig. 2A). The *dpn-GFP*-marked abdominal NSCs showed weak but distinct abdA expression compared to other cells in this region at early L2 (28-32 hr ALH), early L3 (42-48 hr ALH), and Mid L3 (66-70 hr ALH) stages (Fig. 2L-N’). We also examined the expression of Antennapedia (Antp) and Ultrabithorax (Ubx) proteins in NSCs. They are expressed in the anterior and posterior thoracic regions of VNC. We found that Antp and Ubx were both weakly expressed in NSCs, with Ubx levels being lower than those of Antp. (Supplementary Fig. 2. I-J’).

To further validate the expression of Hox genes in NSCs, we analyzed available single-cell RNA sequencing (scRNA-seq) data (Dillon et al., 2022). They have sequenced single cells from larval CNS at various developmental time points (1 hr, 24 hr, and 48 hr ALH). We first clustered cells using the standard Seurat package and identified clusters expressing known NSC markers such as *grh*, *mira, dpn*, *CycE*, *wor*, *ase*, and *insc* (Supplementary Fig. 2. F-H). Several cells within the NSC cluster were positive for the four Hox genes. Our analysis revealed expression of *Antp*, *abdA*, and *AbdB* in several cells in the NSC cluster of larval VNC across different developmental time points (Fig. 2 L-N’, O-Q, and Supplementary Fig. 2 I-J’). *Ubx* was also expressed in several NSCs at 1 hr, but the number was reduced at 48 hr ALH.

The difference between our data and previous reports on abdA expression could be attributed to the enhanced sensitivity of our analytical techniques. Since the expression levels of Hox genes in NSCs are lower than in other cells, as demonstrated by immunostaining and single-cell RNA sequencing (scRNA-seq) data, detection may have been challenging. The presence of Hox genes in NSCs suggests a potential post-embryonic role for these cells. Given that elevated levels of abdA are known to induce apoptosis and eliminate NSCs, our findings indicate that the sustained low expression of abdA in abdominal NSCs during the early larval stage is crucial for regulating the NSC behavior during larval development.

### The canonical apoptotic pathway does not regulate the size of abdominal NSCs

During development, embryonic and larval abdominal NSCs are eliminated via apoptosis (Arya et al., 2015; Bello et al., 2003; White et al., 1994). The cell death program begins during the late embryonic stage, dramatically reducing the number of abdominal NSCs (Abrams et al., 1993; Arya et al., 2015a; Peterson et al., 2002; White et al., 1994). The three surviving abdominal NSCs per larval VNC hemi-segment, vm (ventro-medial, 5-2), vl (5-3 ventro-lateral), and dl (dorso-lateral, 3-5), undergo apoptotic elimination at approximately 72 hr (Bello et al., 2003; Bossing et al., 1996; Schmidt et al., 1997a; Truman and Bate, 1988). Thus, despite the presence of several NSCs in the thoracic VNC, the late abdominal VNC is devoid of NSCs. It is important to note that the Hox gene *abdA* is required for NSC apoptosis in both embryonic and larval stages (Arya et al., 2015; Bello et al., 2003).

Loss of cell volume or cell shrinkage is a characteristic of apoptotic cell death. In the *Drosophila* CNS, mushroom body NSCs in the central brain also show retarded growth and a reduction in size before their elimination by apoptosis and autophagy (Siegrist et al., 2010). Thus, we hypothesized that the reduction in NSC size could be linked to the activation of cell death pathways. To determine whether apoptotic players regulate abdominal NSC size, we blocked apoptosis using different members of the cell death pathway in the hierarchy (Fig. 3A). First, we examined the role of *rpr*, *hid*, and *grim* genes, the most upstream activators of apoptotic signaling, which are transcriptionally activated by death signals (Peterson et al., 2002; White et al., 1994), in regulating the size of abdominal NSCs. To deplete *reaper*, *hid*, and *grim* expression in NSCs, we used a microRNA (*UAS-RHG-miRNA*) (Siegrist et al., 2010) that targets all three apoptotic genes. Reduction in the expression of these cell death activators in NSCs (*insc>mCD8GFP;RHG-miRNA*) permitted the continuous survival and proliferation of abdominal NSCs until the third instar stage due to a failure to undergo apoptosis (Tan et al., 2011, Bello et al., 2003). Significantly, the rescued abdominal NSCs were as small as the wild type in the mid-L3 larvae (Fig. 3B, C, D). To further confirm these results, we used homozygous deletion of the MM3 regulatory genomic region, which controls the expression of the reaper, grim, and sickle pro-apoptotic genes (Tan et al., 2011) (Fig. 3A). In agreement with the above results, deletion of the MM3 regulatory region resulted in the survival of abdominal NSCs until the late larval stages. However, their nuclei were as small as in wild-type (Supplementary Fig. 3 B, C). Taken together, our data indicate that apoptotic signaling does not regulate the size of abdominal NSCs.

**Figure 3.**
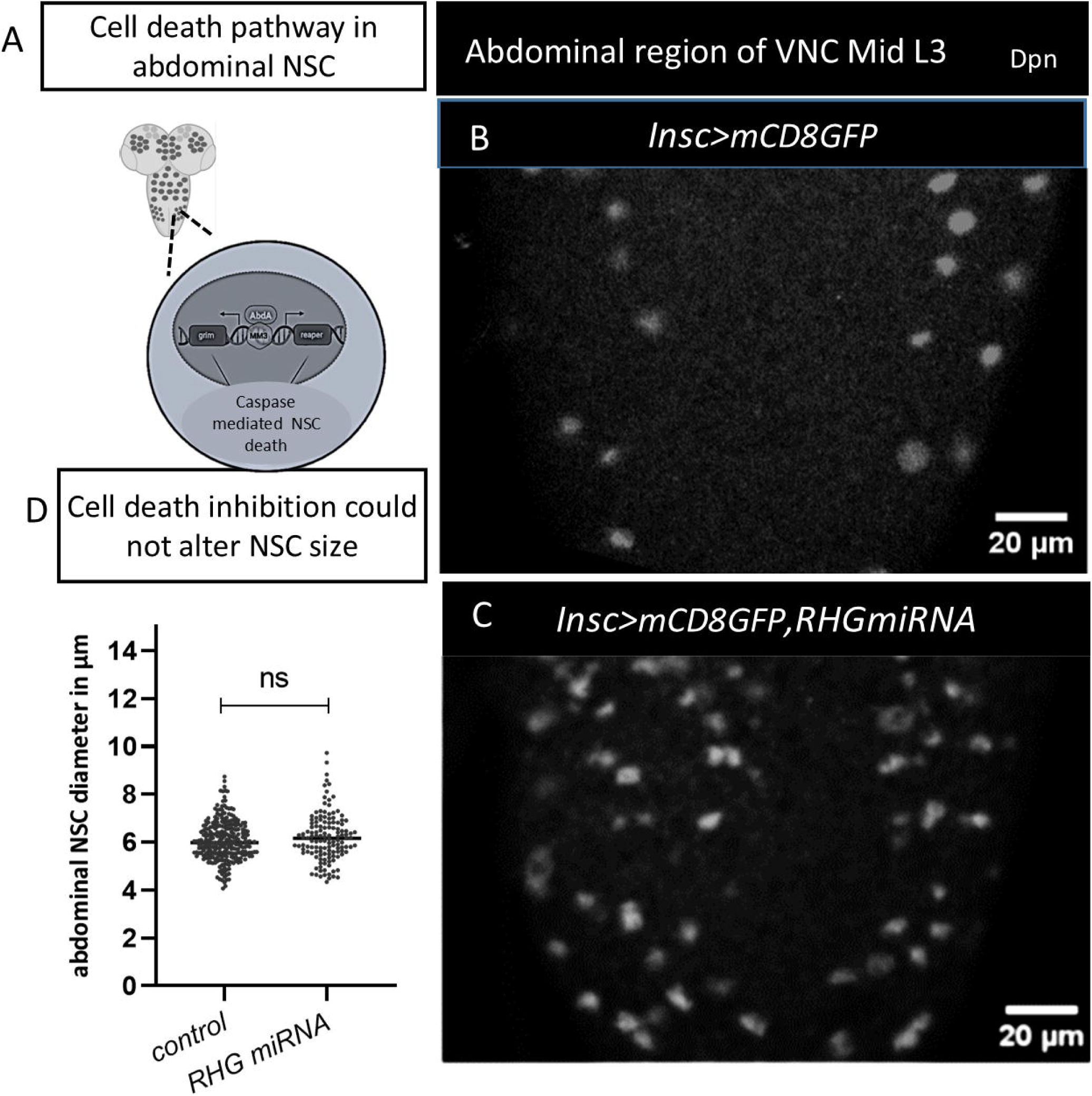
The canonical cell death pathway does not regulate the size of the abdominal NSCs. (A) A model depicting abdominal NSCs apoptotic regulation by abdA. (B-C) nuclei of NSCs (marked with Dpn) in the abdominal region of Mid L3 in control(*insc>mCD8GFP*) and rescued in RHGmiRNA (*insc>mCD8GFP,RHGmiRNA*) are quantified in (D) for the size showing no significant difference. The statistical evaluation of significance based on an unpaired t-test is marked with asterisks, ****P<0.0001. Scale bar: 20 µm. More than 5 CNS were analyzed in the control and *RHGmiRNA*.

### *abdA* regulates the timely entry of abdominal NSCs into mitosis and retards their cycle rate

Proper regulation of the rate and duration of NSC proliferation is essential for maintaining the volume, shape, and function of the CNS. Our research indicates that the *abdA* regulates the size of abdominal NSCs, keeping them smaller than thoracic NSCs. Notably, while both thoracic and abdominal NSCs enter the S phase at approximately 25 hours ALH), abdominal NSCs do not become mitotically active until around 50 hours ALH(Taylor and Truman, 1992; Truman and Bate, 1988). Since abdominal NSCs proliferate more slowly and produce significantly fewer progeny than thoracic NSCs (Truman and Bate, 1988), we investigated whether *abdA* influences the delayed entry of abdominal NSCs into mitosis as well as the rate of their division. Following *abdA* knockdown, several abdominal NSCs in mid-L3 stage produced more progeny than control cells of the same age (see Fig. 2E, F for GFP-positive counts per NSC). For an NSC to generate a larger lineage, either the duration of its proliferation needs to increase, or the proliferation rate must accelerate. We examined both, whether the timing and duration of mitosis are affected in *abdA-*depleted NSCs.

Initially, we assessed the timing of NSC mitotic entry. As previously discussed, abdominal NSCs have a longer G2 phase compared to thoracic NSCs (Truman & Bate, 1988). We further confirmed it with the help of FUCCI tool (Supplementary Fig.4 A-D) (Zielke et al., 2014). Abdominal NSCs do not enter mitosis and remain in G2 until after 50 hours ALH, despite progressing to the S phase between 24 −36 hours ALH (Truman & Bate, 1988; Fig. 4A). In the *abdA* knockdown VNC, abdominal NSCs had already produced 2-6 progeny at 48-52 hours ALH (early L3), while control cells showed no signs of proliferation, indicating that *abdA*-depleted NSCs began dividing much earlier (see Fig. 4A, D).

**Figure 4.**
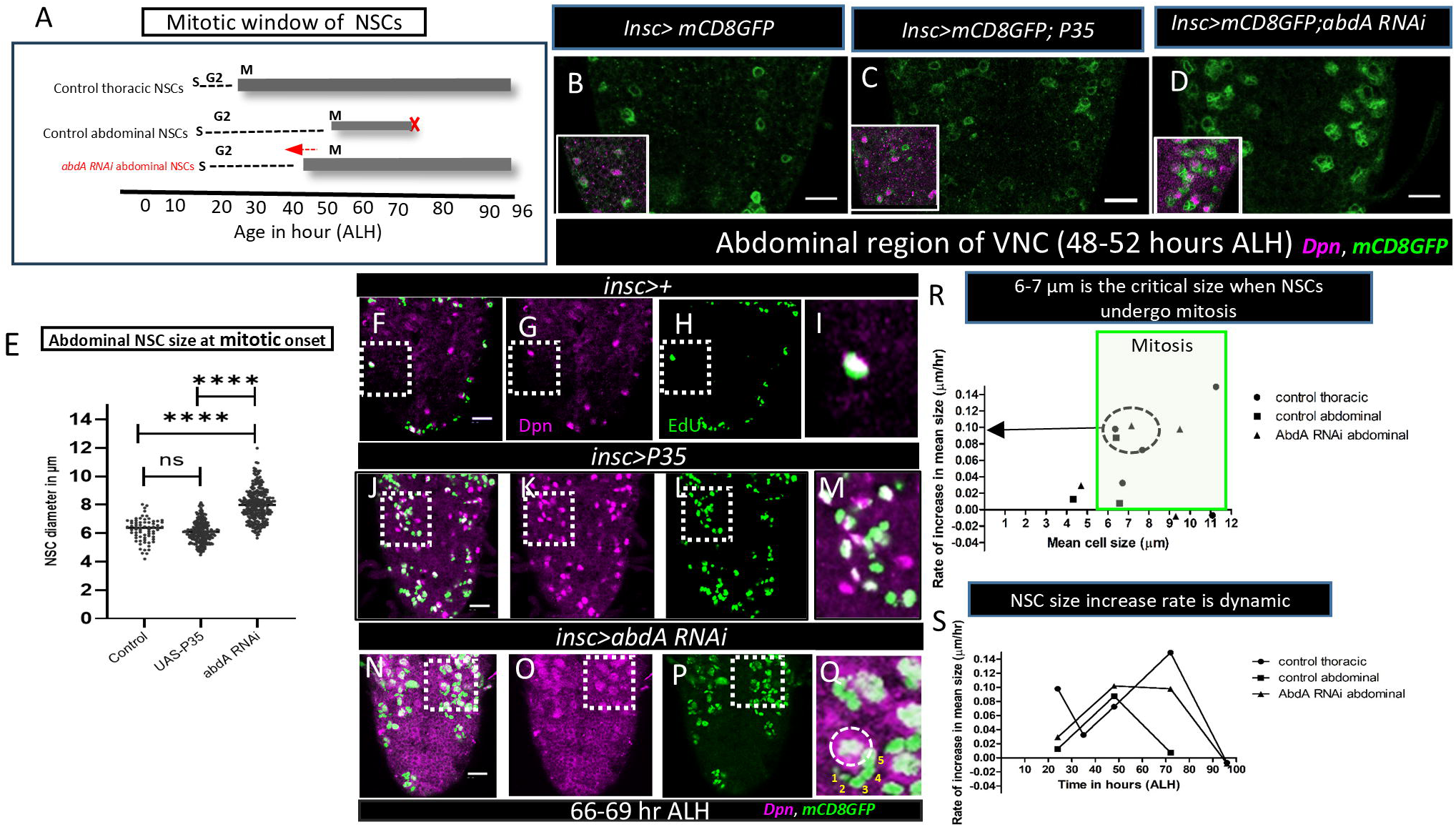
*abdA* regulates the timely entry of abdominal NSCs into mitosis and retards their cycle rate. (A) Schematic showing the duration of active cycling of thoracic and abdominal NSCs, gray bar depicting the active proliferation window in wild type, and arrow showing the shift of the NSC proliferation to an earlier window upon abdA-depletion (S,G2, M are various cell cycle phases). (B-D) 48-52ALH abdominal NSCs in control (B,C *insc>mCD8GFP* and *insc>mCD8GFP;P35* respectively) were quiescent and did not produce progeny, whereas in *abdA-*knockdown VNC (D, *insc>mCD8GFP; abdA-RNAi*), NSCs entered mitosis and formed progeny; insets show NSCs (Dpn, magenta) and NSC and their progeny (mCD8GFP, green). (E) Quantification of the NSC size for the datasets shown in B-D. (F-Q) EdU (green) incorporation in abdominal NSCs (megenta) in control group (*insc>+* and *insc>P35)* and *abdA RNAi* (*insc> abdA-RNAi)*, at 66 to 69 hr ALH; dotted squares are shown in zoom (I,M,Q) with 1-5 numbers in (Q) indicating more progeny per NSC upon *abdA*-knockdown. (R) Graph showing the critical size of NSCs for entering mitosis with their different rate of increase in size. (S) Graph showing the dynamic pattern of rate of NSC size increase in control and *abdA* knockdown during larval developmental stages. The statistical evaluation of significance based on an unpaired t-test is marked with asterisks, ****P<0.0001. Scale bar: 20 µm. More than 5 CNS were observed in each case.

To ensure that the progeny observed in the *abdA* knockdown VNC were not present during the embryonic stage, we examined the larval VNC at an earlier developmental window of 21-26 hours and found no progeny at this stage (Fig. 2B). This indicates that most of the abdominal NSCs undergo early quiescence exit and become mitotically active between the mid-second larval stage (L2) or just before the onset of the early third larval stage (L3). Since inhibiting apoptosis by expressing P35, the broad-spectrum caspase inhibitor, does not affect the mitotic window of the abdominal NSCs (Fig. 4A, B), we eliminated the possibility that the inhibition of cell death is responsible for this early entry into mitosis (Fig. 4 compares C and D).

We also evaluated the size of the abdominal NSCs in control, P35, and *abdA* knockdown conditions at 48-52 hours ALH. The abdominal NSCs in the *abdA-*knockdown VNC were approximately 1.3 times larger than those in the control and P35-expressing conditions (Fig. 4E). Therefore, we propose that the larger abdominal NSCs enter mitosis earlier following the loss of abdA.

Thoracic NSCs divide faster than abdominal NSCs (Truman and Bate, 1988). Specifically, a thoracic NSC completes its division in approximately 55 minutes, while an abdominal NSC takes more than 2 hours to complete a single cell division (Truman and Bate, 1988). To investigate whether the *abdA* also influences the rate of cell cycle progression, we conducted an EdU incorporation assay for 3 hours during the mid-third larval stage (66-69 hours ALH), a period during which abdominal NSCs actively proliferate (Fig. 4A). Since the progeny of NSCs remain associated with their parent NSC, forming a cluster, we compared the number of EdU-positive cells per cluster to determine how many progeny cells are generated by a single NSC. In the control group of abdominal NSCs, we observed a maximum of 3 EdU-positive cells (one NSC and two progeny) (Fig. 4F-M). In contrast, the *abdA-*depleted NSCs produced 4 to 5 progeny in several instances (Fig. 4N-Q). Since many rescued NSCs incorporate EdU and produce progeny that are often in close proximity to one another, it was challenging to identify individual NSCs and their progeny. Therefore, we examined the overall pattern to assess the maximum number of progeny produced by a single NSC. This led us to question whether the rate of NSC size increase over time could influence the number of progeny produced. Our mathematical analysis of size increase rates versus time indicates that within the mitotic window, the rate of NSC size increase is dynamic, measuring between 0.0-0.16µm/hr for thoracic NSCs and 0.00-0.1 µm/hr for abdominal NSCs. Both types of NSCs reach approximately 6 µm in size before entering mitosis (Fig. 4R,S). Interestingly, *abdA-*depleted NSCs initially increase in size at the same rate as control abdominal NSCs but maintain that size for a longer duration before experiencing a decline (Fig. 4Q). This data strongly suggests that the larger size of *abdA-*depleted NSCs leads to their earlier entry into mitosis, enabling them to produce progeny at a faster rate (Fig. 4 F-Q and R, S). Additionally, the possibility that *abdA* regulates the mitotic rate directly in these NSCs warrants further investigation.

The enlargement of NSC in size is necessary for their entry into the cell cycle (Britton & Edgar, 1998; Chell & Brand, 2010; Yuan et al., 2020). To evaluate the rate of NSC size increase and to determine whether a size threshold is required to enter mitosis, we analyzed this parameter in relation to developmental timing and NSC size in both the thoracic and abdominal regions. Our findings suggest that a cell size threshold is indeed necessary to exit G2 quiescence and initiate the first mitosis. The average size of thoracic and abdominal NSCs at the time of mitotic entry is approximately > 6 µm, suggesting that this is the critical size required for NSCs in the VNC to enter mitosis. Interestingly, altering the levels of *abdA* in the NSCs of the VNC—either through knockdown in abdominal NSCs or ectopic expression in thoracic NSCs—affects the NSC size increase and their subsequent mitotic entry. As previously discussed, when *abdA* is knocked down, abdominal NSCs reach the critical size earlier than 50 hr ALH and enter mitosis before their designated developmental time (Fig.2 A-H, 2J). Conversely, thoracic NSCs that ectopically express *abdA* do not reach the critical size by 31 hours ALH; they remain at 5 µm and are unable to enter mitosis. (Fig.5, details discussed in the next section). Thus, NSCs in the VNC require a minimum critical size for mitotic entry. Together, this finding indicates that the loss of *abdA* not only increases the size of abdominal NSCs and allows them to reach the critical size earlier, but also enhances their division rate, enabling them to generate more progeny.

**Figure 5.**
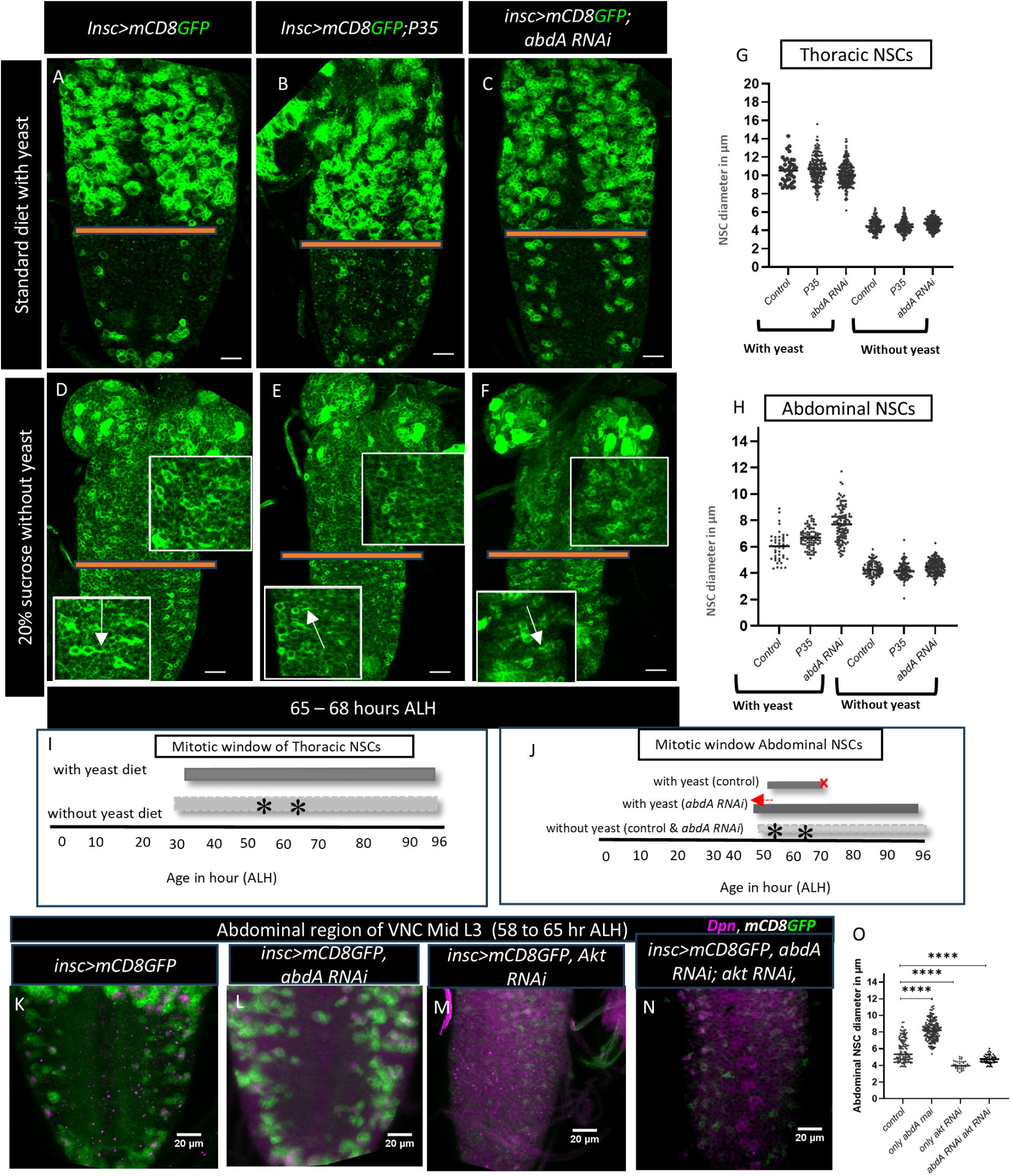
Nutrition is vital for NSC reactivation, and *abdA* helps fine-tune their size increase post-reactivation. (A-C) Well-fed larvae at 65-68 hr ALH thoracic and abdominal NSCs in (A,B) control (*insc>mCD8GFP, insc>mCD8GFP; P35*) and (C) *abdA-*knockdown (*insc:mCD8GFP>abdA RNAi*) showing active proliferation and the presence of GFP-positive progeny cells. (D-F) Under starved conditions, neither controls (D,E) nor NSCs with abdA knockdown (F) showing signs of mitotic entry and quiescence exit. Insets in (D-F) show zoomed-in views and arrow show tail-like projections of the quiescent NSCs. (G, H) Quantification of NSC size data for thoracic and abdominal NSCs presented in A-F. (I,J) Summary diagram showing the active proliferation windows in thoracic and abdominal NSC under fed and starved conditions. The star (*) indicates the time at which we evaluated the proliferation status of NSCs. The arrow showing the NSC proliferation window shifted earlier in *abdA-knockdown cells*. (K-N) abdominal NSC size and their proliferation status in control groups *(insc>mcD8GFP, insc>mCD8GFP, abdA RNAi)* and upon *Akt* knockdown condition with and without *abdAi (insc>Akt RNAi* and *insc> abdA RNAi; Akt RNAi)*, at Mid L3 larvae (58 to 65 hr ALH). (O) Quantification of NSC size data presented in K-N. The statistical evaluation of significance based on an unpaired t-test is marked with asterisks, ****P<0.0001. Scale bar: 20 µm. More than 5 CNS were analyzed in each case.

### Nutrition is vital for NSC reactivation, and *abdA* helps fine-tune their size increase post-reactivation

Several studies have demonstrated that the growth of NSCs is dependent on nutritional signals, which are essential for their size increase and transition into the S phase and mitosis. (Britton & Edgar, 1998; Chell & Brand, 2010; Yuan et al., 2020). When larvae begin to feed actively, circulating dietary amino acids in the hemolymph are detected by fat bodies, which then secrete fat-body-derived mitogens (FBDM) into the hemolymph (Britton & Edgar, 1998). These FBDM stimulate surface glia to release *Drosophila* insulin-like peptides (dILPs), which trigger NSC growth through dlnR/PI3K/Akt signaling (Chell & Brand, 2010; Sousa-Nunes et al., 2011). Under nutrition-restricted conditions, the thoracic NSCs of larvae do not increase in size or enter the cell cycle(Britton & Edgar, 1998; Chell & Brand, 2010; Yuan et al., 2020).

Interestingly, some NSCs, such as the mushroom body and lateral NSCs in the *Drosophila* brain, do not rely on nutrition for proliferation. These particular cells are larger than other NSCs and begin to proliferate early during larval life, even in the absence of nutrition (Britton & Edgar, 1998; Truman & Bate, 1988; Yuan et al., 2020). We aimed to investigate whether the loss of *abdA* allows abdominal NSCs to increase in size independently of nutrient availability. To explore the relationship between nutrition-growth regulating pathways and *abdA* on the size of abdominal NSCs and their proliferation onset window, we selected two developmental time points during larval life: one at which the abdominal NSCs started mitosis, the early L3 (approximately 50 hr ALH), and another near Mid L3 when most of them actively engaged in proliferation (approximately 65 hr ALH) (Fig. 5J). Under nutrition-restricted conditions, neither thoracic nor abdominal NSCs grew or entered the cell cycle in the control, as previously reported (Britton and Edgar, 1998; Chell and Brand, 2010; Yuan et al., 2020) Inhibition of cell death by expressing P35 in NSCs also did not lead to cell cycle entry (Fig. 5D,E, G-J). Similarly, *abdA-*downregulated abdominal NSCs did not increase in size or enter mitosis even at 65 hr ALH (Fig. 5F,G-J). We compared the size of Mid L3 NSCs (65-68 hr ALH) in the sucrose-fed yeast deprived control group and *abdA-*knockdown animals. found that in both cases, the sizes of thoracic and abdominal NSCs were smaller than those in the fed larvae and they also did not enter mitosis (Fig. 5A-H). These smaller NSCs also have a tail-like projection peculiar to quiescent NSCs (arrow in inset of Fig. 5D-F) (Truman & Bate, 1988), These findings indicate that nutrition is critical for initiating the growth of NSCs and subsequently entering mitosis, even for NSCs with depleted *abdA*.

Nutrition is known to act through insulin-dependent PI3K/Akt signaling to reactivate NSC growth, and ectopic activation of PI3K and Akt can initiate growth, allowing NSCs to attain a larger size and proliferate, even in the absence of nutrition (Chell and Brand, 2010). We further examined whether *abdA-*depleted NSCs could increase their size without Akt. We observed that Akt-depleted NSCs did not grow in size or proliferate, even under nutrient-supplemented conditions (Fig.5. K-N). Interestingly, NSCs showed no signs of growth when both *abdA* and *Akt* were depleted (Fig. 5.K-N, O), indicating that activation of nutrient-derived signaling is essential to reactivate NSCs, and *abdA* is required to determine the rate of NSC size increase downstream of this activation.

Our results suggest that the size-related control of *abdA* acts downstream or in parallel to nutritional growth signaling when NSCs are activated and capable of growth. Upon receiving nutrition, both thoracic and abdominal NSCs begin to grow; however, abdominal NSCs exhibit slower growth compared to their thoracic counterparts. We propose that *abdA* in abdominal NSCs limits their size, growth rate and overall proliferative capacity once they are reactivated. In summary, *abdA* is sufficient to restrict the size increase of reactivated abdominal NSCs and delay their entry into mitosis.

### Regulation of neural and gut stem cell size and proliferation by ectopic expression of *abdA*

We demonstrated that *abdA* controls the regulated size increase of abdominal NSCs. To investigate whether *abdA* is also sufficient to regulate the size of other NSCs, we examined whether its ectopic expression could similarly restrict the size of thoracic NSCs. We expressed *abdA* in NSCs from embryonic life (*insc>mCD8GFP,abdA*) and evaluated the thoracic NSCs size. It is important to note that ectopic expression of Hox genes, including *abdA*, triggers the apoptosis of NSCs (Bello et al., 2003). To study the role of *abdA* in growth regulation, we inhibited NSC apoptosis by co-expressing the caspase inhibitor, P35 (*insc>mCD8GFP,abdA;P35*). Interestingly, we noted that thoracic NSCs expressing *abdA* showed growth retardation over time (Fig. 6. A-D I). in the Late L3, the size of the thoracic NSCs in the control group ranged from 8-14µm, while following the expression of *abdA*, their size reduced to 5-12 µm, which is approximately 1.2 times smaller than those in the control (Fig. 6E-H, J). We inferred that *abdA* could instruct a size retardation program in NSCs (Fig. 6K,L).

**Figure 6.**
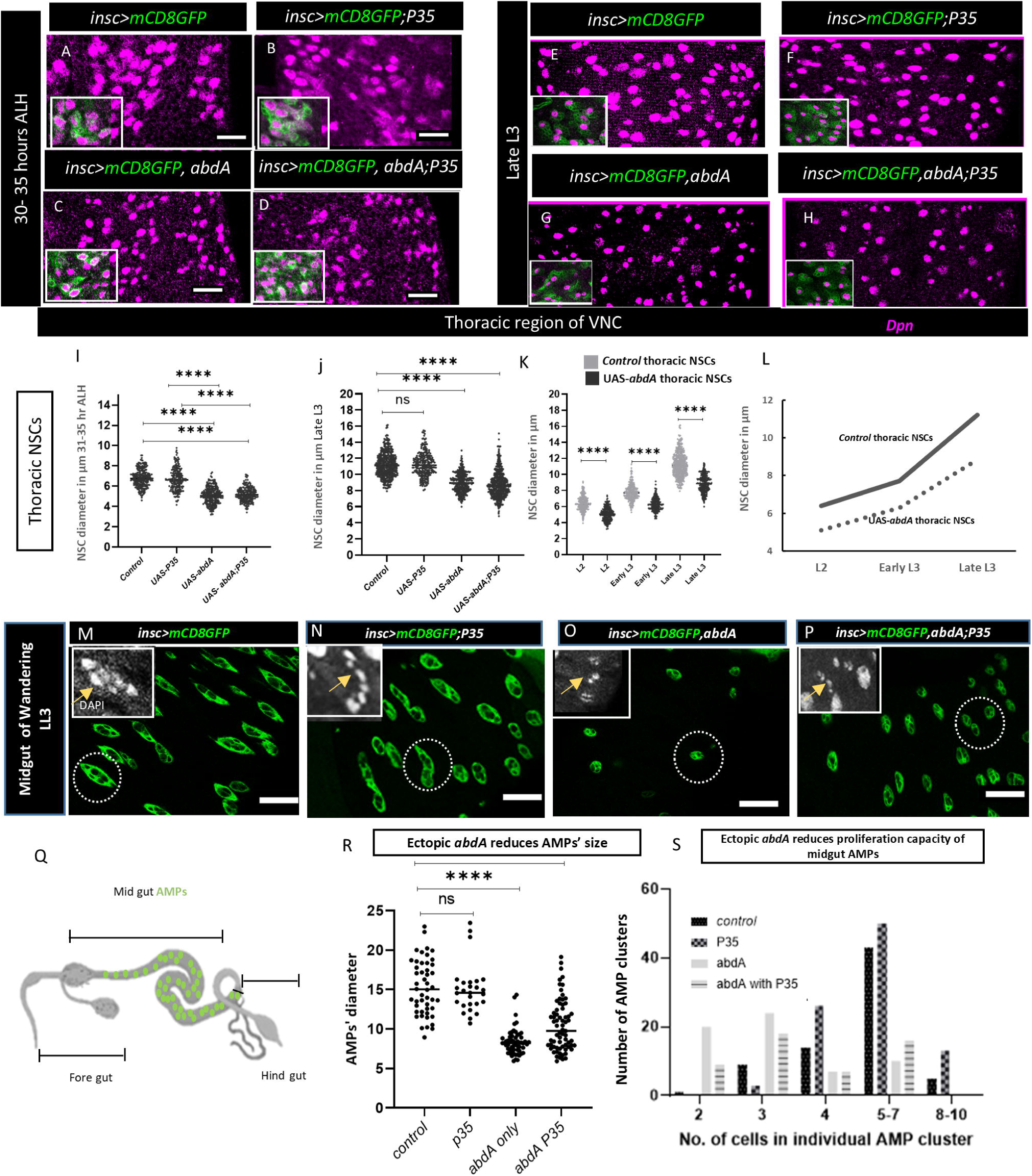
Regulation of neural and gut stem cell size and proliferation by ectopic expression of *abdA*. Thoracic NSC size upon ectopic *abdA* expression evaluated at (A-D) 30-35 hr ALH and (E-H) late L3 larval stages. (A, B, E,F) controls (*insc>mCD8GFP, insc>mCD8GF;P35*), and (C,D,G,H) ectopic *abdA* expression (*insc>mCD8GFP,abdA, and Insc>mCD8GFP,abdA;P35*). Note that the NSCs are smaller in C,D and G,H compared to A,B and E,F respectively. (A,B) controls already have proliferating NSCs at this window (insets show GFP positive lineage) while (C,D) ectopic *abdA* did not enter mitosis at the designated time (insets show no lineage around NSCs). (I,J) Quantification of NSC size of genotypes presented in A-D and E-H respectively. (K, L) Quantification of size of the thoracic NSCs during different larval stages showing ectopic *abdA* retard their size increase. (M-P) Size of *Drosophila* mid gut progenitors AMPs and proliferation capacity reduced upon *abdA* ectopic expression. AMPs are marked with GFP and cell nuclei marked with DAPI (inset, grey), one islet is marked with dotted white circle. (M-N) control (*insc*>mCD8GFP, *insc>mCD8GFP;P35*) and (O-P) ectopic *abdA* expression (*insc>mCD8GFP,abdA, and Insc>mCD8GFP,abdA;P35*). (Q) A model showing different parts of *Drosophila* gut, AMPs are depicted in midgut region (green). (R) quantification of AMP size of genotypes (M-N). (S) number of cells present in a single AMP islet (produced by individual AMP) in mid gut of different genotypes (M-N). The statistical evaluation of significance based on an unpaired t-test is marked with asterisks, ****P<0.0001. Scale bar: 20 µm. More than 5 CNS and gut were analyzed in each case.

Given that ectopic expression of *abdA* diminished the size of thoracic NSCs, we also investigated whether these smaller NSCs entered mitosis at the appropriate time. Typically, thoracic NSCs exit quiescence at 28-30 hr ALH (Truman & Bate, 1988) (Supplementary Fig. 6A). Our findings revealed that ectopic expression of *abdA* not only restricted NSC growth but also delayed their entry into mitosis, causing them to remain non-dividing at 31-35 hours ALH (Fig 6. A-D, G-I). Taken together, we conclude that *abdA* regulates the size and timing of NSC entry into mitosis. The two effects are strongly correlated but could reflect independent activities of *abdA*.

To investigate whether *abdA* influences the size of different types of stem cells, we focused on intestinal stem cell-like progenitor cells known as adult midgut precursors (AMPs) (Fig.6Q). These cells exhibit similarities in their differentiation processes and asymmetric division with NSCs Guo and Ohlstein, 2015; Jiang and Edgar, 2009; Plygawko et al., 2024; Wu et al., 2023; Zhang et al., 2024). AMPs, which originate from endoblasts (the embryonic progenitors of AMPs), begin a process of self-amplification during the larval stage (Green et al., 1993; Jiang and Edgar, 2009; Plygawko et al., 2024).

During the early larval stages, AMPs multiply and disperse to form islets throughout the midgut. As larval development progresses into the later stages, the dividing AMPs become confined to these islets. By the wandering third instar stage, there are approximately eight AMPs per cluster, which can increase to about thirty cells per cluster just after pre-pupa formation (Jiang and Edgar, 2009). Since *abdA* is not expressed in larval AMPs (Supplementary Fig. 6B,C), we used *insc-Gal4* which is also expressed in larval AMPs, to drive ectopic expression of *abdA*, (Pandey and Kumar Roy, 2024). This approach allowed us to examine the effect of *abdA* on AMP’s size and proliferation. Similar to its role in inducing cell death in NSCs, the ectopic expression of *abdA* in AMPs also led to apoptosis, which was partially rescued by the co-expression of P35 (Fig. O-P, R-S.). In the control midgut, the size of AMPs ranged from 9-20 µm, with several islets containing 5-7 cells. However, ectopic expression of *abdA* reduced the AMP size to 5-11µm and resulted in 2-3 cells in several islets (Fig. 6.M-S). Therefore, we conclude that *abdA* regulates the size and proliferation of NSCs and gut AMPs.

## Discussion

### Regulation of the size of NSCs is essential to realize their potential

Our study highlights the functional importance of regulating NSC size in determining their spatial and temporal fate within the developing *Drosophila* CNS. Once born during the embryonic CNS development, NSCs in the thoracic and abdominal regions of the VNC are initially identical in number and organization, dividing to produce neurons and glial cells. (Ito et al., 2013; Knoblich, 2008; li Ming & Song, 2011; Schmidt et al., 1997; Yu et al., 2013, Doe, 1992). Despite all NSCs in various hemi-segments of the embryonic CNS being born in a stereotypical manner and sharing a common identity, later their individual identity and fate, as well as the number and type of progeny they produce, are influenced by their spatial location in the CNS and unique gene expression patterns(Doe, 1992; Doe and Technau, 1993; Hartenstein and Campos-Ortega, 1984; Prokop and Technau, 1991; Sen et al., 2019; Skeath and Thor, 2003; Verma et al., 2019). By the end of embryonic life, most surviving NSCs in the VNC attain a small size and enter a quiescent state (Hartenstein et al., 1987). These quiescent NSCs reactivated by increasing their size during larval life, and resume the cell cycle to produce neurons in adult flies (Truman & Bate, 1988, prokop and Technau 1991). Notably, NSCs in the brain and thoracic region of the VNC continue to produce neurons into the pupal stages, whereas abdominal NSCs generate neurons only until the larval stage (Bello et al., 2003; Truman & Bate, 1988). Interestingly, NSC size is closely correlated with exit from quiescence and may determine the timing of exit. We found that when NSCs in the VNC reach a size of approximately 6μm, they exit quiescence. During the active division period of larval life, the NSCs continue to grow bigger in size and also after every division the thoracic NSCs regrow and maintain their size, but later during pupal life their size gradually decreases before undergoing symmetric division to form two terminally differentiated cells(Homem et al., 2014, Murange et al 2008). Similarly, the size of mushroom body NSCs in the fly brain also decline before being eliminated by autophagy and apoptosis induction (Siegrist et al., 2010).

We found that abdominal NSCs are the smallest population of NSCs of all types of NSCs present in the *Drosophila* CNS, and that the *abdA* Hox gene regulates their size (Figure 7 model). Their size increase is notably slow, with a prolonged G2 phase, leading to a delayed entry into mitosis. We show that abdA expresses in all abdominal NSCs from the early larval stage and retards the increase in their size and delays their entry into mitosis compared to thoracic NSCs. (Truman and Bate, 1988). Remarkably, while thoracic NSCs complete a cell division cycle in about an hour, abdominal NSCs take at least twice as long (Homem et al., 2013; Truman and Bate, 1988). We observed that *abdA* plays a crucial role in regulating the cell cycle rate of NSCs. The persistent presence of *abdA* in these cells prevents them from growing as large as their thoracic counterparts, resulting in slower cycling and fewer progeny. The delayed mitosis entry and also slower rate of division might be due to restricted size increase of abdominal NSCs. Thus, the size of NSCs may have a vital role in controlling neurogenesis in the abdominal region of larval VNC. The size of a stem cell may be pivotal in determining the optimal timing for neuron production and the number of neurons generated to innervate a specific tissue.

**Figure 7:**
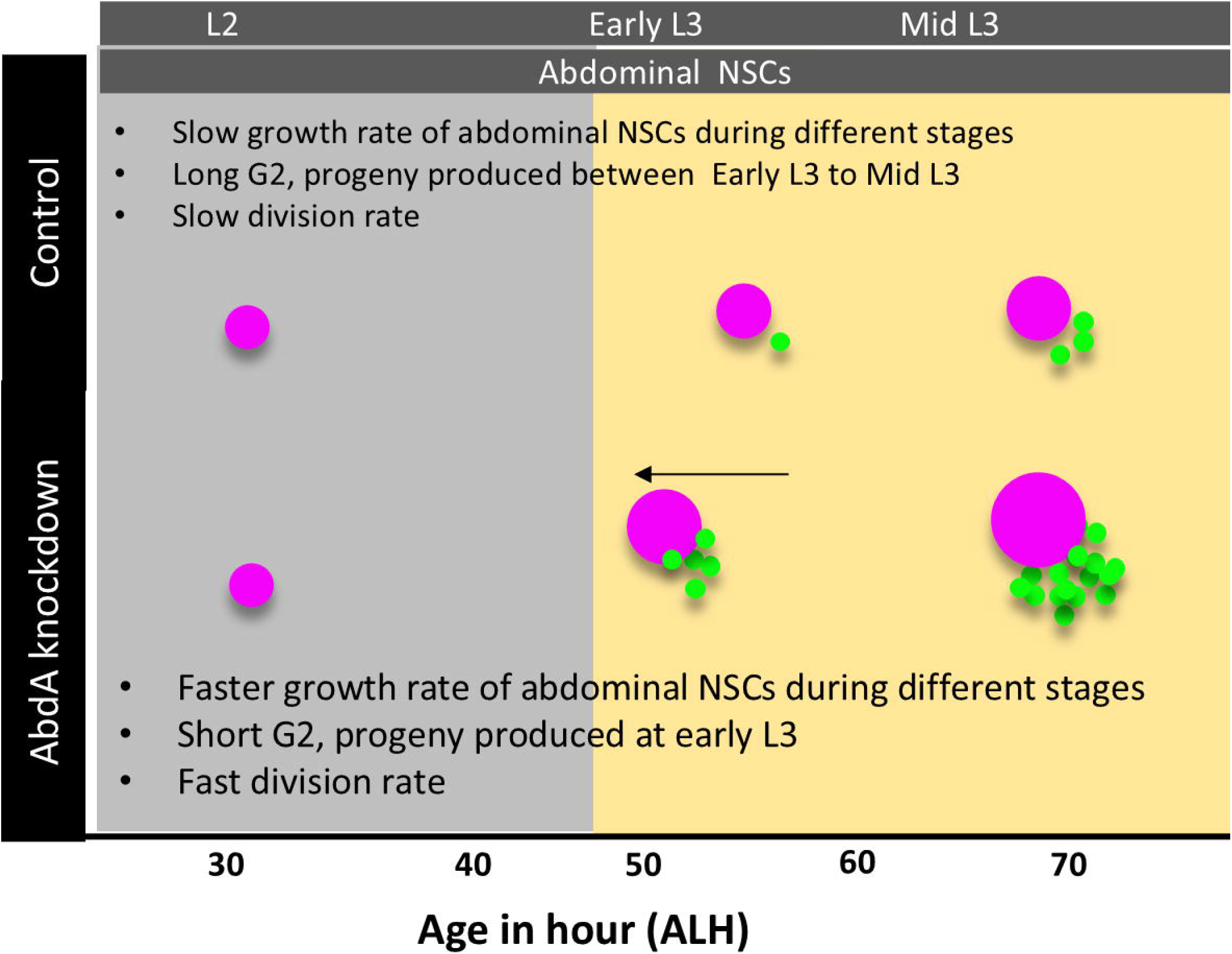
A model depicting change in size of abdominal NSCs and shift in their proliferation window upon *abdA* knockdown.

The regulation of NSC size and mitotic entry involves a complex interplay of signaling, including the insulin, PI3K/Akt/TOR, and Hippo pathways. Insulin signaling plays a vital role in breaking the quiescent state of most type I NSCs (Britton & Edgar, 1998; Chell & Brand, 2010; Ding et al., 2016; Sousa-Nunes et al., 2011). With the availability of circulating amino acids in the nervous system, the PI3K/Akt/TOR pathways become active, thereby removing inhibitory Hippo signaling and initiating NSC growth (Britton & Edgar, 1998; Chell & Brand, 2010; Ding et al., 2016; Sousa-Nunes et al., 2011). Thus, these growth pathways tightly control the switch between the non-dividing state and the proliferative state of NSCs. The availability of nutrients reactivates NSCs, rendering them competent to increase their size and enter mitosis. We noted that the size retardation signal and mitotic entry which are guided by *abdA* only functions when NSCs achieve competence to grow in the presence of a nutritional signal. In a nutrition-deprived state, *abdA* knockdown failed to initiate the size increase of abdominal NSCs and also could not start mitosis even in later stages, indicating that *abdA* may acts downstream of nutrition-mediated activation of NSCs but might fine-tune the downstream signaling to achieve the appropriate size to initiate proliferation of abdominal NSCs. These findings suggest that nutritional signals are responsible for initiating the size increase of all type-I NSCs and that the presence of abdA in abdominal NSCs acts as a growth retardation signal to maintain their required size and mitotic rate. This specific role of *abdA* in NSC growth regulation provides a deeper understanding of the complex mechanisms involved in regulating the fate of NSCs.

### The Hox gene *abdA* is an important fate determinant of abdominal NSCs

During development, Hox genes, including *abdA*, play crucial roles as regulators of specific signaling pathways. They operate in a tissue-specific manner to determine the correct body patterning, including tissue formation at an appropriate location. (Lewis, 1978; Mann & Morata, 2000; Marin et al., 2012; Miller et al., 2001; Rogulja-Ortmann & Technau, 2008). In the *Drosophila* nervous system, *abdA* is a significant determinant of spatiotemporal fate. It is expressed in the central abdominal region of embryonic and larval nervous system (Arya and White, 2015; Lewis, 1978; Sánchez-Herrero et al., 1985). *abdA* is well known for its role in regulating the apoptosis of abdominal NSCs during embryonic and larval life via the canonical cell death pathway (Arya and White, 2015; Bello et al., 2003;Khandelwal et al., 2017). Previously, it was shown that larval abdominal NSCs do not express abdA, and only act as a pulse at a specific time to induce apoptotic signaling (Bello et al., 2003, Prokop and technau 1998). It has been suggested that expression of Hox genes such as *abdA* and *Ubx* in NSCs may create a stable memory imprint during their early expression in embryonic life, which stably regulates the fate, including proliferation, of NSCs even in their absence (Prokop et al., 1998). In contrast, our study revealed that NSCs express *abdA* and other Hox genes, such as *Antp* and *Ubx*, from the very early larval stage, although their expression levels are low. The expression of *abdA* helps control the appropriate size of these cells, regulating their entry into mitosis and their potential for proliferation.

To form a functional CNS, tissue growth is coordinated at multiple levels. In this study, we found that *abdA* regulates the size and mitotic potential of NSCs, determining their capacity to form neurons. In *Drosophila*, post-embryonic NSCs in the larval CNS regain their activity and enter the cell cycle to develop into a functional adult CNS (Prokop and Technau, 1991). Intriguingly, not all NSCs in the CNS start proliferating simultaneously. Instead, they follow a temporal pattern along the anterior-posterior axis to enter mitosis (Truman and Bate, 1988). At approximately 30 hr ALH, the thoracic NSCs entered S phase and started proliferating. It is important to note that the abdominal NSCs enter into S phase at the same time as thoracic NSCs, but they start proliferation 20 hr later, at approximately 50 hr ALH (Taylor and Truman, 1992; Truman and Bate, 1988, Prokop and technau 1991). Therefore, it is important to understand the factors that delay the entry of NSCs into mitosis. Our data strongly suggest that the slow growth of these abdominal NSCs due to abdA expression does not allow them to increase their size and become mitotically active earlier and divide as fast as their thoracic neighbours.

## Conclusion

Our study underscores the critical function of the Hox gene *abdA* in modulating the size of abdominal NSCs and determining their mitotic potential. From the commencement of larval development and the availability of nutrients, both thoracic and abdominal NSCs become reactivated and progressively enter mitosis. However, the rate of increase in the size of abdominal NSCs consistently lags behind that of thoracic NSCs, resulting in their persistent smaller size compared to thoracic NSCs. We propose that *abdA* decelerates the growth rate of abdominal NSCs by prolonging the G2 phase of the cell cycle and regulates their capacity to produce only a limited number of neural cells within the appropriate temporal window. The absence of *abdA* expression in NSCs permits them to increase in size earlier, enter mitosis sooner, and initiate neurogenesis in an ectopic window, potentially disrupting normal organismal development. Thus*, abdA* slows the growth rate of abdominal NSCs to extend the non-dividing period and spatially and temporally control neurogenesis in this region. *abdA* is also known to regulate the timely removal of these NSCs from the nervous system. This finding holds significant implications for our understanding of neurogenesis and role of Hox gene *abdA*, as it provides insights into the regulatory mechanisms and their importance in governing the behavior and fate of NSCs for subsequent neurogenesis.

## Materials and Methods

### Fly stocks

*Drosophila melanogaster* was reared at 25°C ±1and crosses were performed at 25 °C in an incubator on a standard food medium containing sugar, agar, maize powder, and yeast. Appropriate fly crosses were set up following standard method to obtain the progeny of desired genotypes (Yadav et al., 2024). The following fly stocks were used in the experiment: wild-type Oregon R +, *inscutable-Gal4,* also known as *1407-Gal4* (a generous gift from Dr. J. A. Knoblich, Vienna, Austria) recombined with UAS-mCD8GFP (BL-5137), *abdA-RNAi* (BL-35644, v106155), *dpn-GFP* (BL-59755) *UAS-P35* (BL-5073), *UAS-abdA* (BL-912), *UAS-miRHG* (obtained from Iswar K Hariharan), *MM3* (from Tan et al., 2011), *tubGal80ts* (7108), *Worniu-Gal4* (56553), *UAS-Akt RNAi* (BL-33615), *UAS-FUCCI* (55122).

We generated second chromosome transgenic fly for *abdA RNAi* (DNA construct# dsRNA-GLV21008, kindly provided by DRSC/TRiP Functional Genomics Resources & DRSC-BTRR, Boston USA) and injected in *Drosophila* Attp40 fly stocks (BL-25709) by Fly Facility at C-CAMP, Bengaluru, Karnataka, India. Click or tap here to enter text.

### Immunostaining, confocal microscopy, and documentation

Larvae F1 progeny of different ages from L1 (17-18 hr ALH), L2 (21-26 hr ALH), Early L3 (48-52 hr), Mid L3 (66-73 hr) and Late L3 (LL3, 80-84 hr ALH) obtained from set crosses of specific genotypes, larvae were selected by choosing GFP reporter and CNSs were dissected in Phosphate buffer solution (PBS 1X containing NaCl, KCl, Na2HPO4, KH2PO4, pH-7.4), fixed in 4% Paraformaldehyde for 30 min., rinsed in 0.1% PBST, (1X PBS, 0.1% Triton X-100), then added in blocking solution (0.1% Triton X-100, 0.1% BSA, 10% FSC, 0.1% deoxycholate, 0.02% Thiomersol) for 1 hr at room temperature. The tissues were incubated with the following primary antibodies: rat anti-Dpn (1:150,195173 and abcam), chicken anti-GFP (1:1000,A10262, Invitrogen), and kept at 4°C for two consecutive overnight incubations, anti-abdA (c-11) (sc-390990, Santa Cruz). The following day, tissues were rinsed thrice with 0.1% PBST (15 min each), incubated with 1 a200 dilution of appropriate secondary antibodies, anti-rat 546 (A11081 and Alexa Fluor, Invitrogen), anti-chicken 488 (A11039 and Alexa Fluor, Invitrogen), and anti-mouse 647 (A21235 and Alexa Fluor, Invitrogen) overnight at 4 °C, and incubated for 2 h at room temperature. Following incubation with secondary antibodies, the samples were washed three times with 0.1% PBST and mounted with DABCO (D27802; Sigma-Aldrich) for further analyses.

Images were acquired using a Leica SP8 STED confocal microscopy facility at the CDC, BHU,India and Zeiss LSM-510 meta, Department of Zoology BHU, India, Nikon A1SiR confocal at MGH USA, Images were assembled using Adobe Photoshop and MS Power Point.Images, Models in figures 1K, 3.A, 6Q were created using BioRender software.

#### Gal80 experiment

For Gal4 suppression by Gal80 expression, we used a temperature-sensitive form of Gal80. The Gal4 activity was suppressed by keeping the egg at 18 °C till they hatch and become 0-4 hr larvae. To activate Gal4 expression, 0-4 hr larvae were shifted to 29 °C with standard food. At the age of 62-66 hr ALH (Mid L3), the CNS were dissected and immunostaining was done using Dpn to mark the NSCs. Following genotype combinations were used for fly crosses: *Gal80^ts^::Wor-Gal4/+ and Gal80^ts^::Wor-Gal4> abdA-RNAi*

#### EdU labeling

The 5-ethynyl-2’-deoxyuridine (EdU) assay was performed using a click it-488 (C10337 and Click-it, Invitrogen) by incubating the dissected CNS for 3 hr. After incubation, the CNS were fixed for 20 min in 4% formaldehyde, washed twice with 3% BSA in PBS, and permeabilized with 0.5% Triton X-100 in PBS twice for 20 min. Incorporated EdU was detected by Click-iT fluorescent dye azide reaction in accordance with the manufacturer’s instructions protocol given in the kit. After immunostaining with Dpn antibody, tissues were stained with DAPI (#10236276001 Sigma-aldrich)

### Image analysis using Fiji and Graph Pad prism

All images were quantified using the Fiji/Image J software (NIH, USA). To measure the NSC size, a freehand tool was used to mark the length and width of the NSC. As the NSC shape was not fully round, two perpendicular lines were drawn along the center of the NSC at its widest point, and their average was considered to be the NSC diameter. Size was measured using the analysis measure option in Fiji.

Graphs were created using GraphPad Prism9 and Ms Excel. Statistical analysis was performed using GraphPad Prism9, where two-tailed unpaired t-tests were used to evaluate statistic<0.05(****P<0.0001, ***P<0.001, **P<0.01, *P<0.05) considered statistically significant, and P>0.05 considered nonsignificant.

Violin plots were constructed using the GraphPad Prism9. The spread of NSC diameter is shown from the median and quartile ranges of distribution in individual groups.

### Nutritional regimen

Control larval groups were fed on a standard *Drosophila* diet of maize, agar, sucrose, yeast, and propionic acid. Starved larvae were yeast-deprived and fed on 20% sucrose diet (Britton & Edgar, 1998). Sucrose-containing 1x PBS solution-soaked cotton was given to the 20% sucrose containing agar matrix for yeast deprived larvae. After hatching the larvae from the embryo, they were transferred to yeast-free food media.

### Single cell RNA sequencing (scRNA-Seq) analysis

Processed scRNA data sets (Seurat files) at 1 hr, 24 hr, and 48 hr after larval hatching (ALH) were obtained from published data sets (Corrales et al 2022). Matrix, barcode, and feature files were downloaded from GEO (GSE135810). five samples were used - GSM 4030601, GSM 4030603, GSM 4030605, GSM 4030613 and GSM 4030614. All analyses were performed using the Seurat R package version 5.1.0. Samples corresponding to the same time point were merged to create three unified seurat objects corresponding to each time point.

Low quality cells (less than 200 features, less than 10% ribosomal RNA and more than 10 % mitochondrial RNA) were not included in the analyses for each Seurat object. Reads for mitochondrial and ribosomal genes were excluded. Principal Component analysis was done using the top 3000 highly variant features. Elbow plot was computed to decide the number of Principal Components (PCs) to be used for downstream analysis. We used the first 30 PCs to cluster the cells using the default Louvain algorithm. The default resolution was used to cluster the cells. 8331, 7657, and 6380 cells were obtained at 1 hr, 24 hr, and 48 hr, respectively. Clusters were identified using known markers as described by Dillon et al. 2022. The expression of four Hox genes known to be expressed in VNC, *Antp, Ubx, abdA*, and *AbdB* was studied in these clusters.

### Mathematical analysis for rate of NSC size increase

To calculate the rate of mean size increase of cells: Consider the size of cell at time t_0_ as x_0_, and the size at time t_1_ as x_1_; the rate of increase r_1_ from time t_0_ to time t_1_ can be calculated as: r_1_ = (x_1_-x_0_)/(t_1_ – t_0_).

## Acknowledgements

We thank Profs. S. C. Lakhotia, B.C. Mondal, G. K., and Pandey, for critically reviewing the manuscript and Dr. Akanksh Pandey for helping with gut experiments. We thank Ramkrishna Mishra and Vaishali Yadav for their assistance during the experiments. We thank the Bloomington *Drosophila* Stock Center (BDSC), NIH P40OD018537, for providing fly stocks and FlyBase release (FB 2022_03) for information on fly genes. We also thank the Cytogenetic Laboratory, Department of Zoology, Banaras Hindu University for all instrumental facilities.

## Author contributions

All the authors contributed to the study. The work started in KW lab at MGH, USA. P.D. performed the animal experiments, designed the workflow, image acquisition, quantification of acquired data, statistical analysis, and created a figure panel under the supervision of R.A. Data curation and analysis presented in Fig. 2O-Q, Fig. 4F and G and Supplementary Fig. 2D-F were done by S. M. and E.A., data visualization was done by E.A.

---

**Supplementary figure 1.**
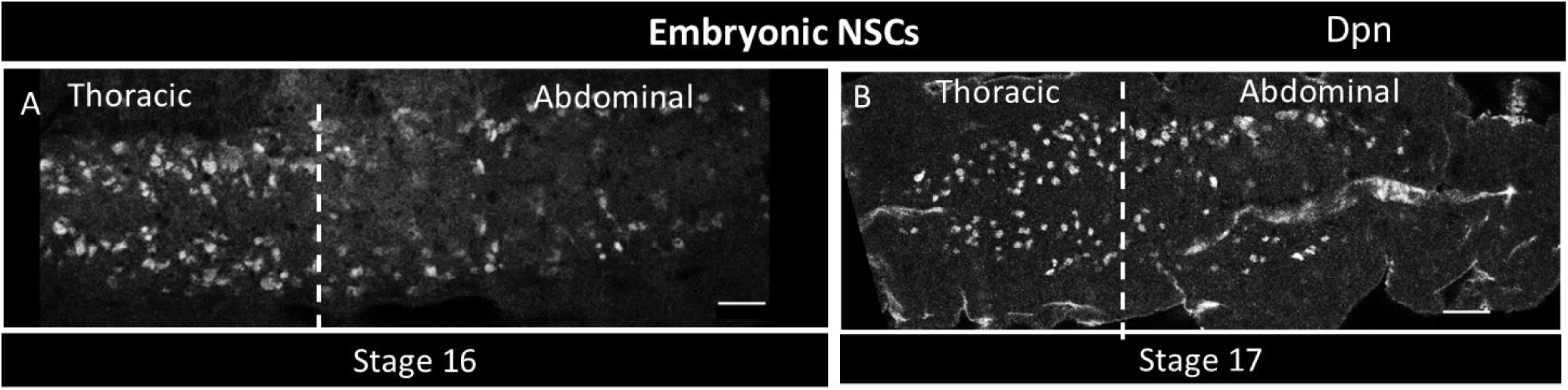
(A,B) Stage 16 and 17 embryos showing NSCs (marked by Dpn) in the thoracic and abdominal VNC (separated by dotted line). Scale bar: 20 µm.

**Supplementary figure 2.**
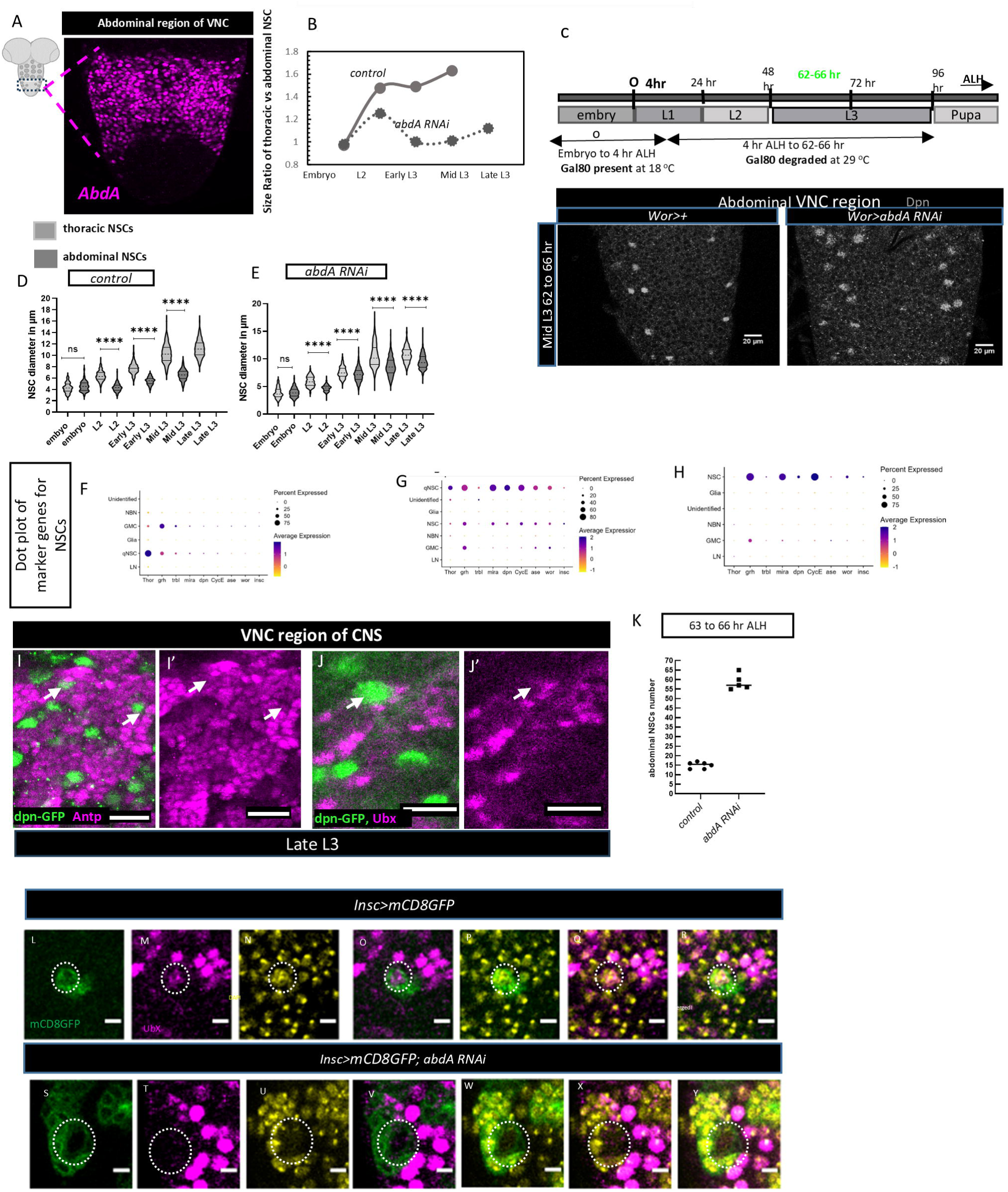
(A) abdA-expressing region in abdominal VNC (B) NSC size ratio in thoracic vs. abdominal regions in control (*insc>mCD8GFP*) and *abdA* knockdown (*insc>mCD8GFP,;abdA-RNAi*). (C) A model depicting temperature restricted expression of *abdA-RNAi* with Gal80ts, control (*wor/CyoGal80ts*) and *abdA* knockdown (*Wor/CyoGal80ts>abdA RNAi) genotypes* (D,E) Showing violin plot forsize of NSCs in thoracic vs abdominal region of VNC in both control *(insc>mcd8GFP)* and upon *abdA* knpckdown (*insc>mcd8GFP; abdA RNAi*). (F-H) Dot plots of marker genes for NSCs at 1 hr, 24 hr and 48 hr. All clusters were identified using markers, as previously described (Dillon et al. 2022). *thor* and *trbl* distinguish quiescent NSCs from other cell types. *grh*, *mira, dpn*, *CycE*, *wor*, *ase*, and *insc* mainly mark type I NSCs and show low expression in quiescent NSCs (qNSCs) at 1h. The qNSC cluster at 1h consists of 466 cells, at 24 hr, it has 55 cells. The type I NSC cluster at 24 hr had 225 cells, and the 48 hr sample had 46 cells in the NSC cluster. (I-J’) Antp and Ubx expression in NSCs. (K) abdominal NSC number in control and *abdA RNAi* (insc>+ and *insc>abdA RNAi*). (L-Y) expression of Ubx in control *(insc>mcd8GFP)* and i knockdown (*insc>mcd8GFP; abdA RNAi*). Scale bar: 20 µm, for Ubx expression scale bar: 5 µm (L-Y)

**Supplementary figure 3.**
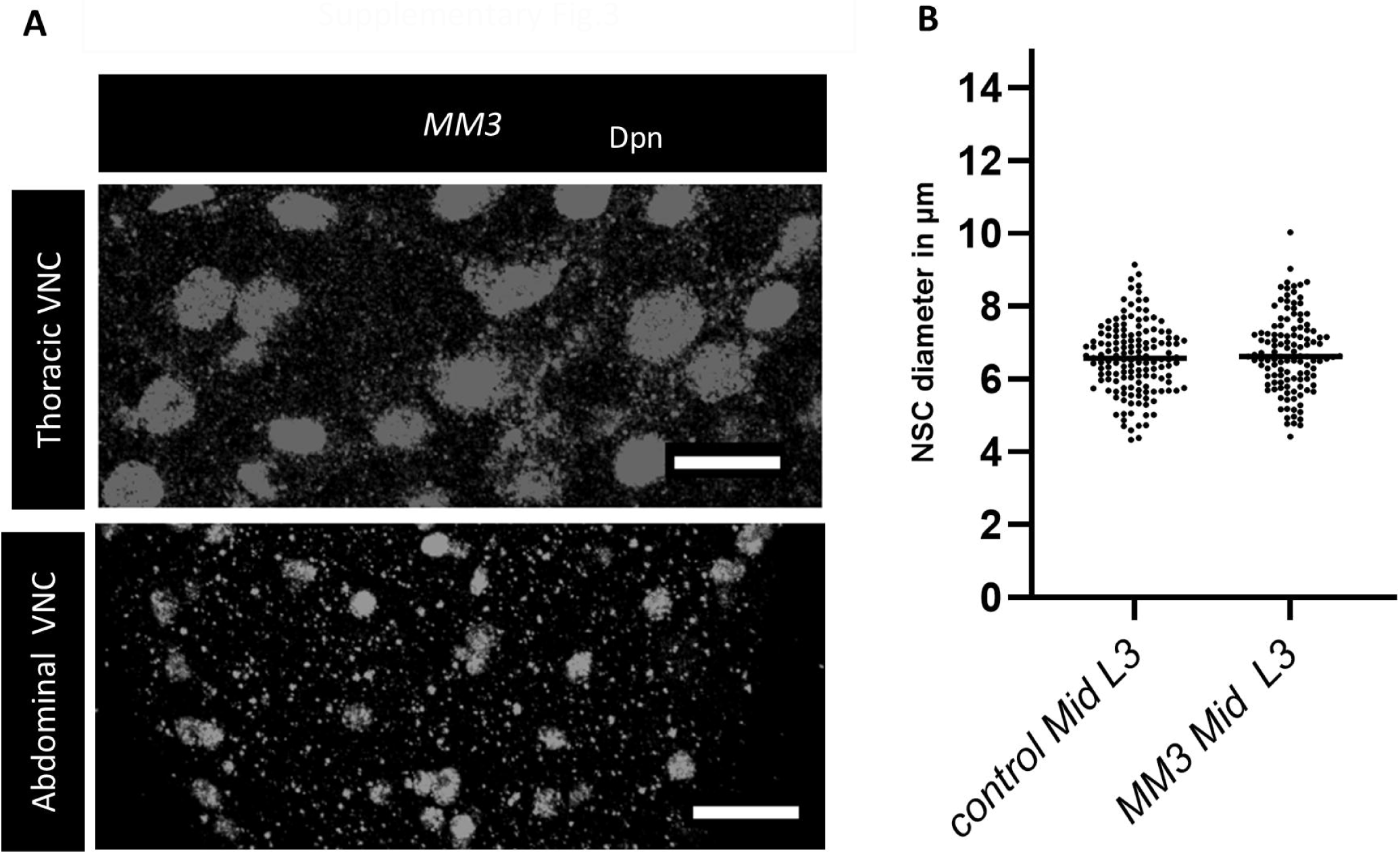
(A) Late L3 VNC of the homozygous *MM3* deletion line, with both thoracic and abdominal NSCs marked by Dpn. (B) *MM3* does not affect the size of the rescued NSCs. The statistical evaluation of significance based on an unpaired t-test is marked with asterisks, ****P<0.0001. Scale bar: 20 µm. N=5.

**Supplementary figure 4:**
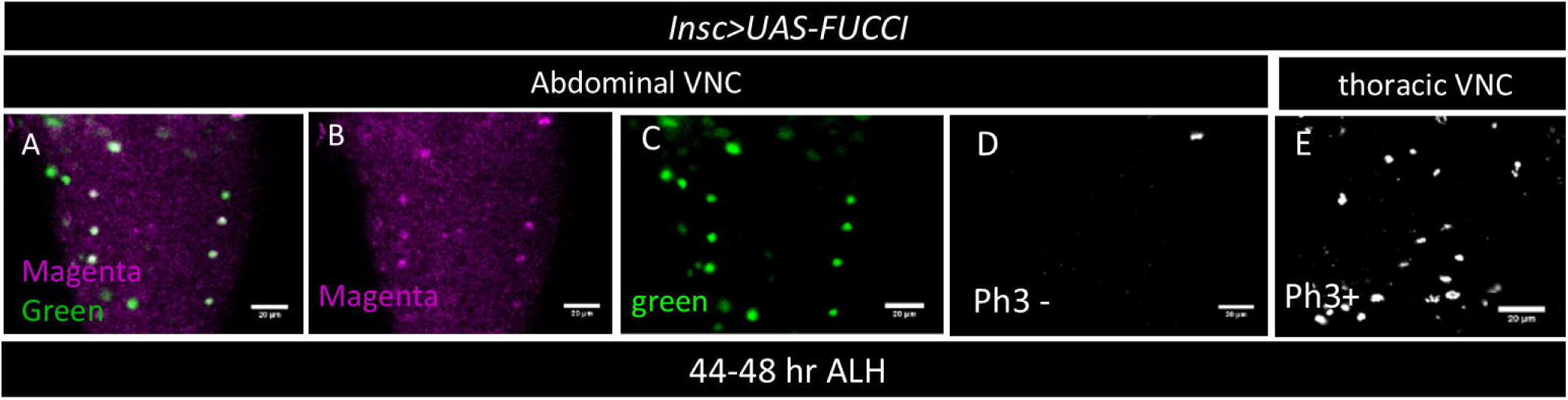
FUCCI tool showing that abdominal NSCs are in G2 phase during Early third (44 to 48 ALH). (A) abdominal NSCs are both magenta and green positive indicating G2 stage (B, C) individual color planes. (D) these NSCs are phosphohistone-3 negative (Ph3^-^) further confirming the G2 stage. (E) thoracic NSCs in the same VNC with Ph3 positive staining showing that ph3 staining worked in the same CNS.

**Supplementary figure 6.**
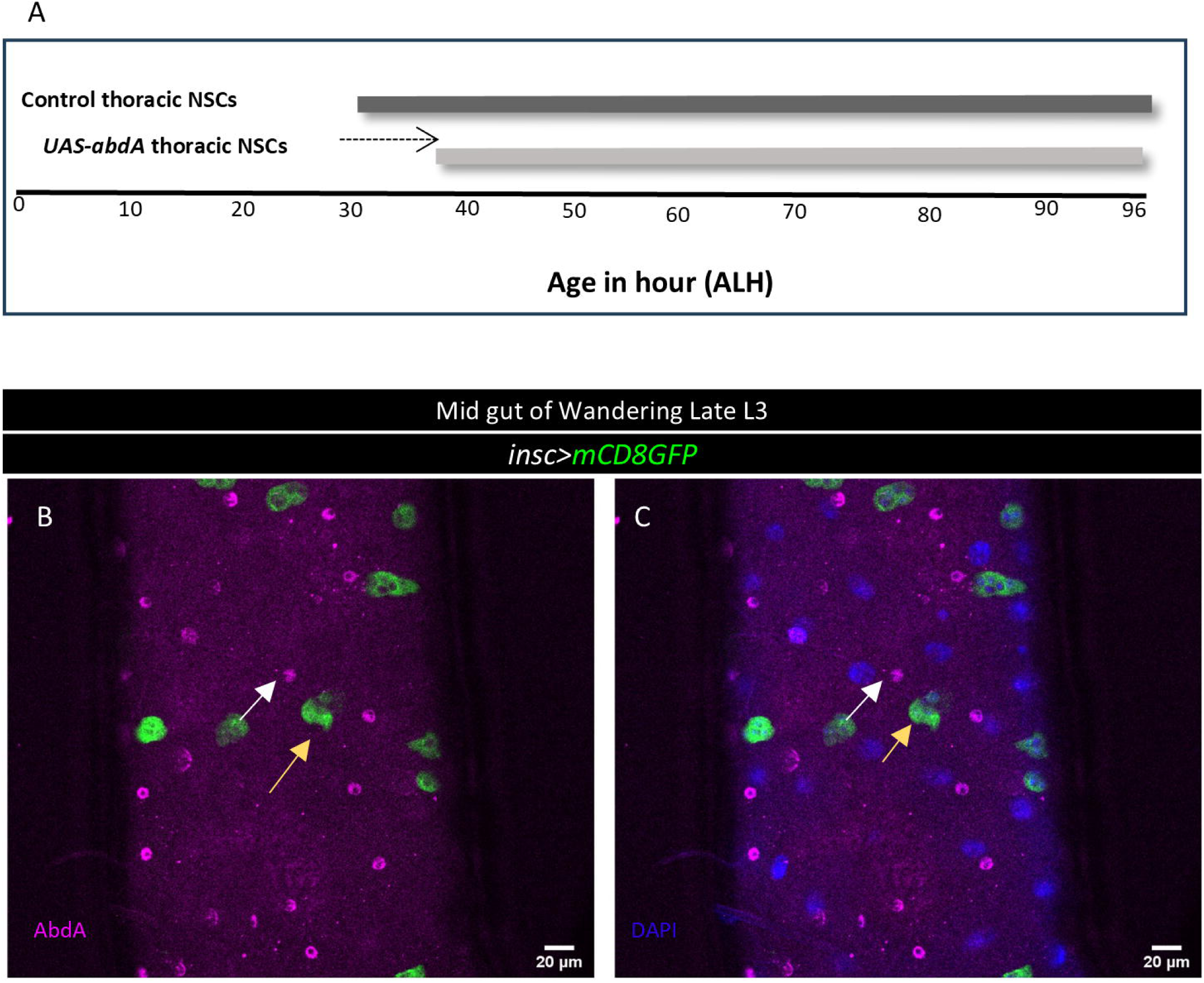
(A) Schematic showing the duration of active cycling of thoracic NSCs, bars depicting the active proliferation window, and arrow showing the delay in the mitotic entry of thoracic NSC upon *abdA* ectopic expression. (B,C) showing *abdA* expresses in the other gut cell types (marked by DAPI) but not in the AMP islets (green).

